# Signal requirement for cortical potential of transplantable human neuroepithelial stem cells

**DOI:** 10.1101/2021.03.27.437311

**Authors:** Balazs V. Varga, Maryam Faiz, Huijuan Yang, Helena Pivonkova, Shangbang Gao, Gabriel Khelifi, Emma Linderoth, Mei Zhen, Samer M. Hussein, Andras Nagy

## Abstract

The cerebral cortex develops from dorsal forebrain neuroepithelial progenitor cells. Initial expansion of the progenitor cell pool is followed by the generation of neurons of all the cortical layers and later, astrocytes and oligodendrocytes. However, the regulatory pathways that control the expansion and maintenance of the neuroepithelial progenitor cell pool are currently unknown. Here we define six basic pathway components that regulate proliferation of cortically specified human neuroepithelial stem cells (cNESCs) *in vitro* without the loss of developmental potential. We show that activation of FGF and inhibition of BMP and ACTIVIN A signalling are required for long-term cNESC proliferation. We also demonstrate that cNESCs preserve dorsal telencephalon-specific potential when GSK3, AKT and nuclear CATENIN-β1 activity are low. Remarkably, regulation of these six pathway components supports the clonal expansion of cNESCs. Moreover, cNESCs differentiate to lower and upper layer cortical neurons both *in vitro* and *in vivo*. Identifying the mechanisms that drive the self-renewal and fate of cNESCs decision of neuroepithelial stem cells is key to developing new stem cell-based therapeutic approaches to treat neurological conditions.

## Introduction

The cerebral cortex develops from the forebrain vesicle of the mammalian neuroepithelium. Forebrain specification of the anterior neuroepithelium happens shortly after neural commitment of the epiblast (Fuccillo et al., 2004). After this initial specification, dorsal forebrain neuroepithelial cells divide symmetrically to expand the progenitor cell pool (Noctor et al., 2004; Smart, 1973). Upon neural tube closure, cells divide both symmetrically to expand the progenitor cell pool and asymmetrically to produce differentiated cells. During early forebrain development, differentiating neural progenitors initially give rise to glutamatergic neurons of the six cortical layers, each with specific connectivity to other brain regions (Molyneaux et al., 2007). At later stages, progenitor cells stop producing neurons and start producing astrocytes and oligodendrocytes (Qian et al., 2000).

The investigation of neural progenitor cell potency, cortical development or pathogenesis in humans is restricted by ethical and technical boundaries. This limitation has been overcome by the *in vitro* use of human pluripotent stem cells; that can be differentiated to a neuroectodermal fate (Chambers et al., 2009) to provide a source of various neural cell types (Yoon et al., 2014). Recent advancement in *in vitro* differentiation of pluripotent stem cells allows generation of three-dimensional cultures of organoids that resemble the structure of the specific tissue. Both monolayer and organoid based methods were improved to make cultures containing progenitors, cortical neurons and glial cells (Bershteyn et al., 2017; Eiraku et al., 2008; Gaspard et al., 2008; Lancaster et al., 2017; Qi et al., 2017; Shi et al., 2012). The cerebral organoids have been used to investigate evolutionary differences between human, chimpanzee and macaque (Otani et al., 2016; Pollen et al., 2019), trace migration of cells or the growth of axons between brain regions (Bagley et al., 2017; Xiang et al., 2017) or to understand the regulation of radial glia progenitor functions (Bershteyn et al., 2017; Nowakowski et al., 2017). Nevertheless, the molecular mechanisms that regulate the differentiation of dorsal forebrain neuroepithelial cells before neurogenesis remain poorly understood.

Maintenance of multipotent neural progenitor cells in the forebrain requires Fibroblast growth factor receptor activity (Paek et al., 2009). However, upon isolation neuroepithelium cells from the embryonic rodent telencephalon show limited proliferation in FGF alone (Pollard et al., 2006). Human rosette stage neuroepithelial stem cells (NESCs) can be maintained in long-term cultures by using Fibroblast growth factor (FGF) at low oxygen (3%) level (Bilican et al., 2014) or adding Epidermal growth factor (EGF). However, forebrain specification is gradually lost; cultured cells acquire posterior identity, do not express dorsal telencephalic markers EMX2, PAX6 and FOXG1 and do not differentiate to TBR1 positive cortex specific neurons (Falk et al., 2012; Hevner et al., 2001; Koch et al., 2009). Complementation of FGF with undefined components of endothelial cell conditioned media can extend the proliferation capacity of the cells and preserve their early cortical potential transiently, indicating that the cells can self-renew if the precise signals are present (Shen et al., 2004). The developing anterior neural plate, and later the forebrain tissue, expresses various extracellular ligands of developmental signalling pathways of BMP/TGFb, WNT, FGF, SHH, NOTCH. These pathways interact with each other, and depending on the timing and strength of each component, progenitor cell fate can be specified. Interaction of FGF, WNT, and BMP pathways is important for induction of the neuroectoderm (Fuentealba et al., 2007; Guo and Wang, 2009) and later, for neural crest cell specification and migration (Martik and Bronner, 2017). Activity of the WNT pathway is mediated by GSK3 inhibition which acts as a hub protein (Doble and Woodgett, 2003; Patel and Woodgett, 2017) regulating the activity of several target proteins involved in cell cycle regulation, mRNA transcription and protein translation. Conversely it has been shown that β-catenin (CTNNB1) transcriptional activity is low during the specification and maintenance of forebrain identity of the neuroepithelium (Ferrer-Vaquer et al., 2010) and this can be mediated by several inhibitory proteins (TLE2, TCF7l1, TCF3, HESX1, CTNNBIP1) (Andoniadou et al., 2011; Masek et al., 2016; Roth et al., 2010; Satoh et al., 2004). *In vitro* stabilization of AXIN2 via chemical inhibition of TANKYRASE by XAV939 can export elevated levels of CTNNB1 from the nucleus and mimic the function of modulatory proteins in stem cells (Kim et al., 2013) (Huang et al., 2009).

Herein we report the identification of six signalling pathway components that are important for the permanent self-renewal of rosette stage cortically specified neuroepithelial stem cells (cNESCs). In addition to FGF, self-renewal requires the simultaneous inhibition of BMP and ACTIVIN A signalling, reduced GSK3 and AKT activity and low level of β-catenin (CTNNB1) nuclear activity. We achieved this complex regulation with six factors. This combination supports single cell-derived cNESC colony formation and preserves the dorsal forebrain-specific differentiation potential of cNESCs. cNESCs express the dorsal telencephalic markers *FOXG1, PAX6, EMX2* and *OTX1/2*, and give rise to glutamatergic projection neurons of lower and upper cortical layers, as well as astrocytes and oligodendrocytes. We also demonstrate that developmental signalling pathways implicated in neural tissue development, such as EGF, SHH and NOTCH signalling, are not required for the self-renewal of the cNESCs. Our results show that neuroepithelial cells with dorsal forebrain specification can self-renew *in vitro* and maintain their early developmental potential.

## Results

### Six signalling pathways regulate the cNESC specification and proliferation

We established dorsal forebrain-specific neuroepithelial stem cells from human pluripotent stem cells (hPSCs) using chemical inhibitors of TGFβ (SB431542) and BMP (LDN193189) signalling (Figure 1A, S1A-C, Supplementary Table 1). Similar to other neural induction protocols (Chambers et al., 2009), efficient neural conversion of hPSCs was seen by day 8, as indicated by the upregulation of neuroepithelial marker, SOX1 (Supplementary Figure 1B,C), downregulation of pluripotency marker, OCT4 (Supplementary Figure 1A), and maintenance of neural stem cell marker, SOX2, in the majority of cells (>95%) (Supplementary Figure 1A). We confirmed neural specification by examining the relative levels of *OCT4* and *NESTIN*, a neural progenitor cell marker in day 11 differentiated NESCs compared to undifferentiated hPSCs. We observed decreased *OCT4* and increased *NESTIN* gene expression in induced NESCs (Supplementary Figure 1D) that was similar to gene expression levels in human fetal forebrain-derived neural stem cells [CB660; (Sun et al., 2008)]. When we examined forebrain specification in these cells, we found that induced NESCs showed increased expression of *EMX2, PAX6, FOXG1* and *OTX2*, region-specific genes that together mark the dorsal forebrain vesicle (Supplementary Figure 1D), and no change in the expression of ventral forebrain markers *LHX6* and *NKX2.1*, ventral midbrain markers *FOXA2* and *EN1*, and hindbrain marker *HOXB1* (Supplementary Figure 1D). Immunocytochemistry confirmed the expression of SOX1, SOX2, FOXG1, NESTIN and PAX6 (Figure 1B-E) and the absence of NKX2.1 expression in NESCs (data not shown). These results indicate efficient induction of dorsal forebrain-specific NESCs from hPSCs within 8-11 days, which we termed cNESCs.

**Figure 1.**
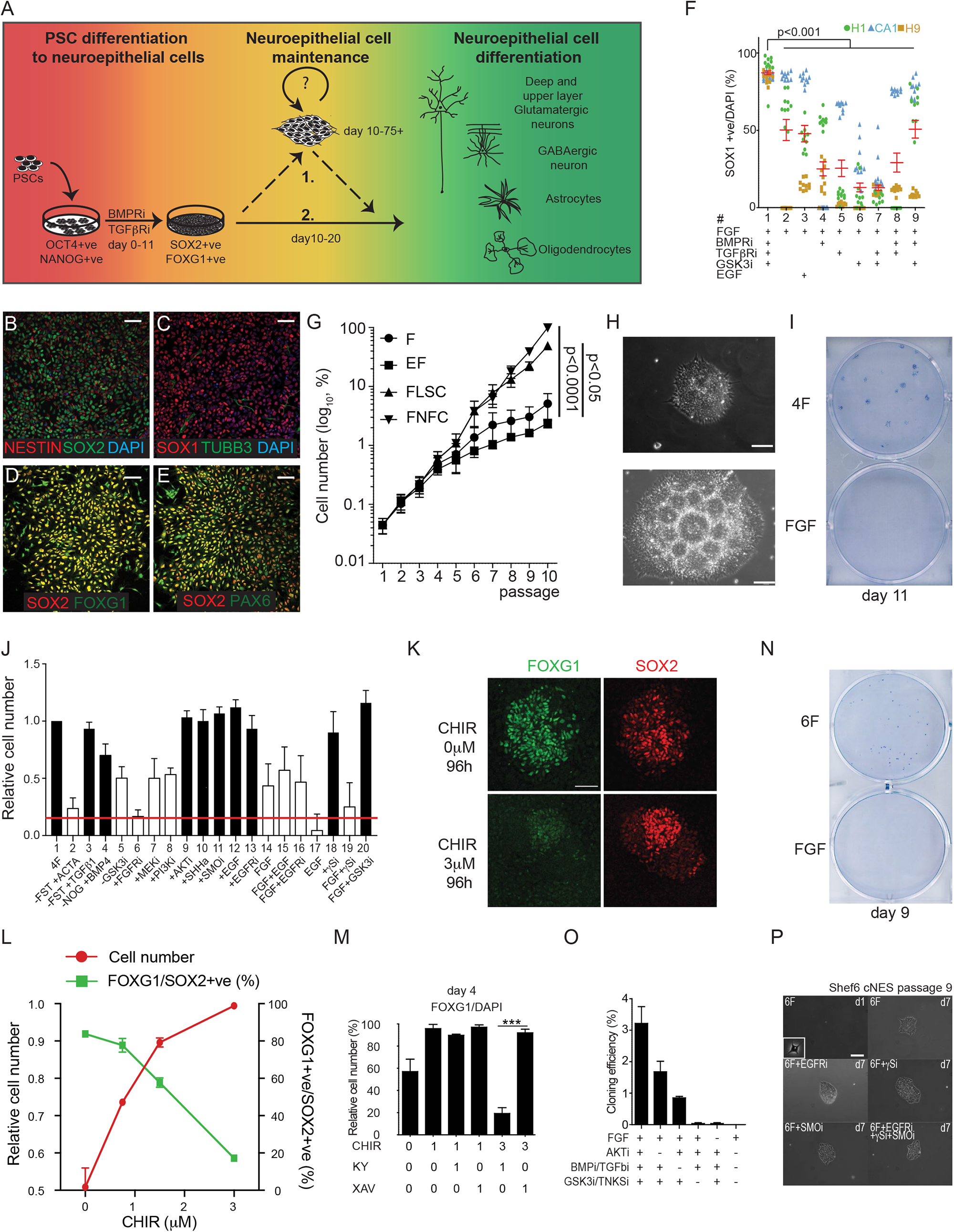
Induction and maintenance of dorsal forebrain neuroepithelial cells. **A:** Neural induction scheme of hPS cells in the presence of TGFbR and BMPR inhibitors. After cortical neural induction NES cells express NESTIN in SOX2 positive cells (**B**) SOX1 in TUBB3 negative cells (**C**), FOXG1 (**D**) and PAX6 (**E**) in SOX2 positive cells. Scalebar: 25μm **F:** SOX1 positive cell ratio in passage 5 cultures of hES (H1, H9, CA1) derived cNES cells treated with various combination of factors (N=3, 10 datapoint per each cell line per group, red bars are mean+SEM, one-way ANOVA, Tukey post-test, p<0.001) **G:** Normalised model of cell number changes of cNES cells (mean+SEM of H1-, H9-, CA1-derived cells, n=3, p<0.001, 2-way ANOVA, Tukey post-tests) over 10 passages (30 days). **H:** Phase contrast image of H1 derived cNES (p36) cells in 4F on day 4 (top) and on day 14 of culture (bottom) forming rosette structures. **I:** Colony formation assay of cNES cells cultured in 4F prior to seeding at 200cells/cm^2^ density. **J:** Quantification of cell number changes after 96-hour treatment of cNES cells with indicated ligands or chemical inhibitors compared to 4F condition (N=3, mean+SEM, 1 way ANOVA, Tukey post-test, white bars are p<0.01, black bars are not significantly different from control). **K:** GSK3 inhibition reduces FOXG1 expression in SOX2 positive NES cells. Scalebar: 25μm **L:** GSK3 inhibition increases cNES cell numbers and decreases FOXG1 positive cell number ratio over a 96-hour treatment in the presence of FGF with or TGFb and BMP inhibition. (N=3, mean+SEM, 1-way ANOVA, Tukey post-test, p<0.001) **M:** Quantification of FOXG1 positive H1 cNES cells after 96 hours culture in 4F media with various concentrations (μM) of GSK3 inhibitor (CHIR) and either XAV939 (XAV) or KY02111 (KY). (N=6, mean+SD, 1-way ANOVA, Tukey post-test, p<0.001) **N:** Colony formation capacity of H1 cNES cells in 6F media or in FGF alone. CNES cells were cultured in 6F media before plating at low density (200 cells/cm2). Colonies were stained after 9 days with cresyl violet. **O:** Quantification of colony formation capacity of H1 cNES cells in 6F, 6F-AKTi, 6F-BMPi/TGFbi, 6F-GSK3i/TNKSi, 6F-FGF, FGF only. (N=3, mean+SD) **P:** CNESCs (Shef6) form colonies in 6F condition in the presence of EGFR, SMO or gS inhibitor or the combination of them. Scalebar: 50μm

### Regulation of BMP, ACTIVIN A and GSK3 regulates long-term maintenance of SOX1-positive cNESCs

Several studies have shown that dorsal forebrain specification is lost during long-term culture of neural stem cells in FGF and EGF (Bilican et al., 2014; Conti et al., 2005; Hitoshi et al., 2002; Sun et al., 2008). Therefore, we asked whether additional extracellular signals were required to maintain FGF-dependent cell proliferation of cNESCs. We investigated modulation of the WNT, BMP/TGFβ, EGF, SHH, NOTCH pathways, which have all been implicated in the regulation of cortical excitatory neuron specification (Hansen et al., 2011).

WNT signals are important positive regulators of forebrain fate (Campbell, 2003) and induce cell proliferation. WNT signal is mediated via the inhibition of GSK3, which can upregulate CTNNB1 and BMP/TGFβ/SMAD signalling (Fuentealba et al., 2007; Guo et al., 2008). Therefore, we tested if FGF in combination with inhibitors of GSK3, BMP and/or TGFβ (GSK3i, BMPRi and TGFβRi, respectively) could maintain the SOX1-positive cNESC population. After neural induction, cNSECs-derived from 3 different hESC lines [H1, H9 (Thomson et al., 1998), CA1 (International Stem Cell et al., 2007)] were propagated for five passages in neural maintenance media with FGF and various combination of inhibitors (Figure 1F). Both FGF alone and FGF and EGF were insufficient to maintain high ratio of SOX1 positive cells. Addition of (4) BMPRi, (5) TGFβRi, (6) GSK3i, (7) GSK3i and TGFβRi and (8) BMPRi and TGFβRi with FGF reduced the ratio of SOX1 positive cells, while addition of (9) FGF, GSK3i and BMPRi showed no change compared to cNESCs in (2) FGF alone or (3) FGF and EGF. Strikingly, we found that neural cells derived from all three hESC lines cultured in FGF and all three inhibitors (termed 4F for 4 factor condition) contained the highest proportion of SOX1-positive cNESCs (mean value 86%; Figure 1F). Conditions 2, 4, 5, 6 and 8 did not support the propagation of at least one of the three different hPSC lines and led to significant cell death, suggesting cell line-dependent variation.

In agreement with other studies (Bilican et al., 2014; Conti et al., 2005; Hitoshi et al., 2002; Sun et al., 2008), we found that cNESCs showed limited proliferation in FGF alone. To determine whether 4F could improve the long-term proliferation of SOX1-positive cells we compared the growth of cNESCs in: 1) FGF alone, 2) EGF and FGF, 3) 4F with chemical inhibitors (FGF, LDN193189 (BMPRi), SB431542 (TGFβRi), CHIR99021 (GSK3i), abbreviated as FLSC) or 4) 4F with protein antagonists (FGF, NOGGIN (BMPi) FOLLISTATIN (ACTIVINi), and CHIR99021 (GSK3i), abbreviated as FNFC. As expected, after six passages, cNESCs in FGF alone and FGF and EGF had a lower proliferation rate than in previous passages. In contrast, cNESCs in both FLSC and FNFC showed sustained proliferation and supported the formation of neuroepithelial colonies (Figure 1G,H). To confirm that the expansion of cNESCs in 4F was not associated with any specific chromosomal abnormalities we examined the karyotype of cNESCs maintained for over 100 days *in vitro* in two independent cell lines and found no chromosomal aberrations (Supplementary Figure 1E).

Next, we tested if 4F was able to support the proliferation of a single cNESC in the absence of additional signals that could be produced by adjacent cells. We plated cNESCs previously cultured in 4F at clonal cell density in either 4F or FGF alone, without supporting cells or conditioned media. cNESCs in FGF alone rarely formed colonies (1 in 3000 cells, N=3) both at low and high oxygen levels (3% and 20%; Figure 1I, S1F). In contrast, single cNESCs in 4F supported single cell derived colony formation at both low and high oxygen levels at an efficiency of around 1%, demonstrating that four pathways can maintain the neuroepithelial specification of a single cNESC (Figure 1I, S1F).

4F did not include developmental factors previously shown to support neural progenitor cell maintenance *in vitro*, such as EGF, NOTCH ligands or SHH (Elkabetz et al., 2008; Sun et al., 2008).

First, we tested their role in the proliferation of cNESCs as we had observed a decrease in the proliferation of cNESCs in FGF or FGF with EGF (Figure 1J bars 14,15). Although cNESCs can activate all three receptors (Supplementary Figure 1H-K), the inhibition of these developmental signalling pathways had no effect on the proliferation of cNESCs in the presence of the 4F (Figure 1J).

Our signalling screening showed that changing the activity of any factor in the 4F condition affects cell proliferation even when the other three components are present (Figure 1J). Activation of ACTIVIN A signalling, removal of GSK3 inhibitor or inhibition of the FGF receptor by PD173074 caused reduced cell proliferation (Figure 1J). FGF signalling appears to be mediated by both the MAPK (bar 7) and PI3K (bar 8) axis, as inhibition of either pathway significantly reduced cNESC numbers (Figure 1O). Our assay also showed that AKT(bar 9) inhibition did not interfere with the cell proliferation in 4F, suggesting that high activity of AKT is not required for PI3K signalling in cNESC maintenance (Figure 1J).

In summary, we determined that FGF signalling through the MAPK and PI3K pathways and inhibition of GSK3 and SMAD signalling are required for cNESCs maintenance, while additional developmental signals like NOTCH, EGF and SHH are not required.

### Six factors maintain dorsal fate and proliferation of cNESCs

Since GSK3 inhibition improved cNESC proliferation in FGF alone or in the 4F (Figure 1J) but early anterior neuroepithelium specification requires low CTNNB1 activity (Ferrer-Vaquer et al., 2010). We tested whether forebrain specification is maintained in the presence of GSK3 inhibition. Gradually increasing GSK3 inhibition (from 0 to 3 μM) in cNESCs caused an increase in the proliferation rate of cNESCs and an immediate decrease in the number of FOXG1-positive cNESCs (Figure 1K, L, S1L). With increasing levels of GSK3 inhibiton, the expression of canonical WNT/CTNNB1 target genes *AXIN2* and *LEF1 increased, EMX2, FOXG1* and *OTX2* mRNA levels decreased, and *PAX6* levels did not change (Supplementary Figure 1M). Together, these results suggest that low levels of both GSK3 and CTNNB1 activity are needed to maintain dorsal forebrain specification in cNESCs.

We aimed to establish the conditions that support cNESC proliferation while maintaining dorsal forebrain specification by regulating the WNT/CTNNB1 activity by lowering the GSK3 activity in combination with low CTNNB1 transcriptional activity. We tested two different chemical inhibitors of CTNNB1 signalling, XAV939 and KYO2111 (Supplementary Figure 1N). XAV939 inhibits TANKYRASE activity, which leads to an increase in AXIN2 levels and reduces CTNNB1 level in the nucleus. (Huang et al., 2009). KY02111 was shown to inhibit CTNNB1 function downstream of GSK3 during cardiomyocyte specification (Supplementary Figure 1N) (Minami et al., 2012). High GSK3 inhibition (3-6 μM) rapidly induced mRNA expression of target genes, LEF1 and EMX2 (Supplementary Figure 1O). In comparison to GSK3 inhibition alone GSK3 inhibition together with TANKYRASE inhibition by XAV939 reduced the transcription of both CTNNB1 target genes to control levels while the addition of KY02111 had no effect.

Next, we tested whether dual inhibition of GSK3 and CTNNB1 transcriptional activity could maintain FOXG1-expression in the cNES cells. Moderate GSK3 inhibition (1μM) resulted the maintenance of a high ratio of FOXG1 and SOX2 double positive cNESCs in the presence or absence XAV939 or KY02111. However, high GSK3 inhibition (3μM) could only sustain high ratio of FOXG1 positive cells in the presence of XAV939 (Figure 1M).

It has previously been reported that neural progenitor proliferation and survival in frog and mouse embryos are regulated by sub-cellular localization of FOXG1 and FOXO proteins via AKT-mediated phosphorylation (Brunet et al., 1999; Regad et al., 2007). We found that FGF signalling activates both MAPK and PI3K/AKT pathways in cNESCs. This was demonstrated by the phosphorylation of ERK and FOXO1/P70S6K and by the reduction in FOXO1 and P70S6K phosphorylation following inhibition of AKT kinase activity (Supplementary Figure 1Q). As we found that AKT inhibition does not interfere with cNES cell maintenance (Figure 1J), we inhibited AKT to enhance FOXG1 function. cNESCs cultured in 4F with TANKYRASE and AKT inhibitor (termed 6F for 6 factor condition) sustained the expression of FOXG1 in cNESCs. Next, we tested whether 6F could support the formation of single cell derived colonies. cNESCs cultured in 6F supported 3 per cent of the cells to form colonies (Figure 1N, O). Removal of either AKT or BMP and TGFβ inhibitors resulted in colony formation, albeit at a reduced level (Figure 1O). Similarly to the 4F condition, inhibition of NOTCH, EGFR or SMO receptors also resulted in colony formation (Figure 1P, S1R). Single cNES cell derived colonies in 6F maintained expression of dorsal forebrain progenitor cell specific genes (Supplementary Figure 1S). However, when single cell derived clones were cultured in 6F media without the TANKYRASE inhibitor (XAV939), an increase in CTNNB1 activity resulted in the increased expression of *AXIN2, LEF1, PAX6* and *NGN1* genes and reduced the level of *FOXG1, OTX2, DLL3, DLL1* and *ASCL1* mRNA (Supplementary Figure 1S) compared to cNESCs in 6F. These results suggest that cNESCs require a balanced regulation of GSK3, CTNNB1 and AKT activity to maintain their dorsal telencephalic fate and that the regulation of these pathways supports the formation of single cell derived colonies.

### 6F cNESCs are bona fide early cortical cells

We wanted to determine whether cNESCs in 6F would maintain the expression of cortical markers FOXG1, OTX1/2, PAX6 during extended culture. cNESCs cultured in 6F for 75 days (15 passages or 50 population doublings) maintained expression of FOXG1 in all five hPSC lines tested (Supplementary Table 2). No decrease in the ratio of FOXG1/SOX2 positive cells (Figure 2B) was observed. Moreover, the expression of high levels of OTX1/2 (caudo-lateral cortex) or PAX6 (fronto-dorsal cortex) was maintained in the SOX2 positive cNESCs from all hPSC lines (5/5; Supplementary Figure 2A, B).

**Figure 2.**
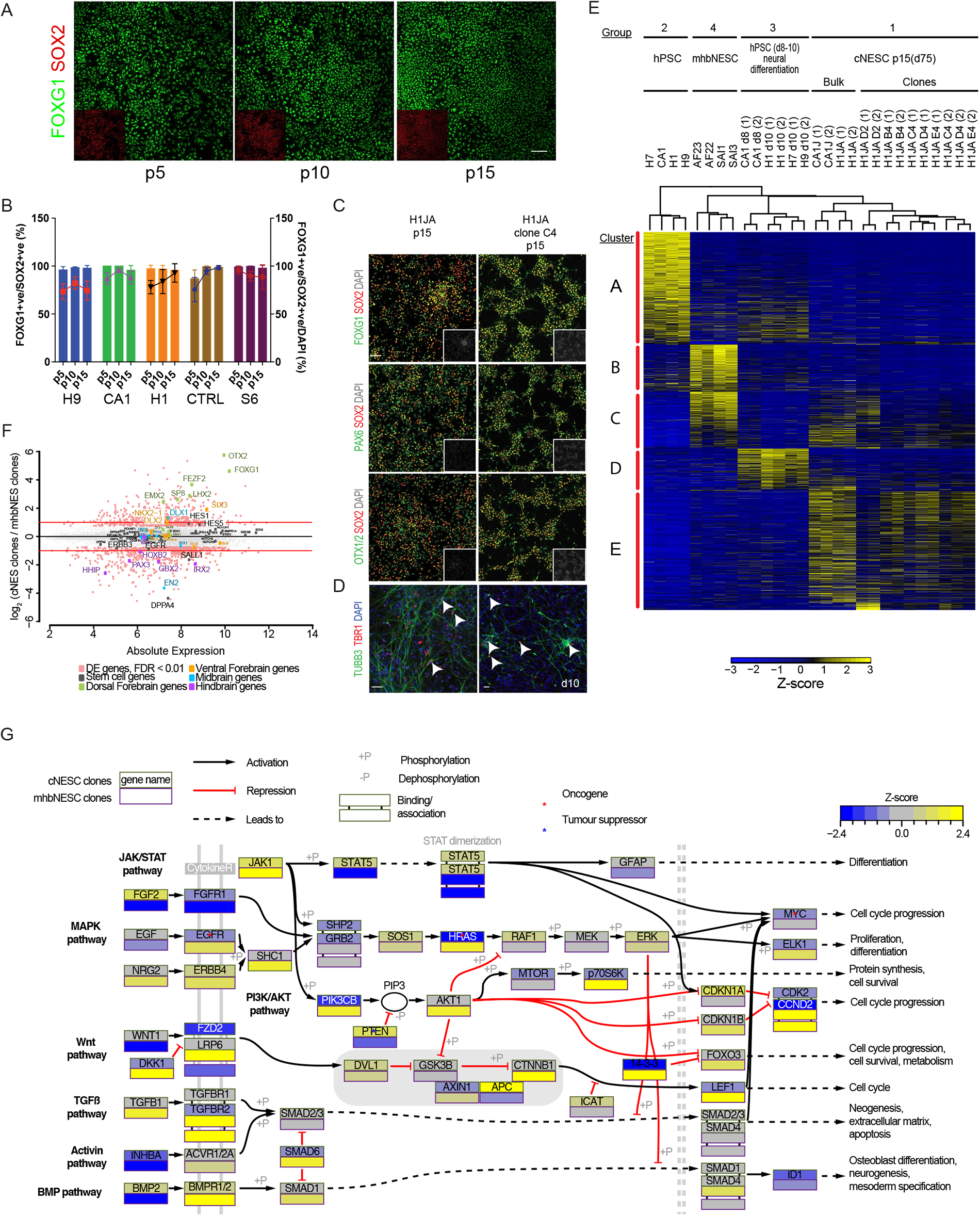
Sustained cortical gene expression in cNES cells. **A:** Immunofluorescent labelling of FOXG1 in H1JA cNES cells cultured in 6F condition from multiple passages. Scalebar:25 μm **B:** Quantification of immunofluorescent labelling of FOXG1 and SOX2 in five independent hPSC derived cNES cell lines from multiple passages. Bars show percentage of FOXG1 positive NES cells, lines show percentage of FOXG1 and SOX2 positive cells of all cells. (Mean±SD) **C:** Single cell derived clones of cNES cells H1JA were cultured in 6F media. Expression of dorsal forebrain markers FOXG1, OTX1/2 and PAX6 was co-labelled with SOX2 neural marker by immunofluorescence in bulk cultures (left) and a clone of H1JA cNES cells (right, H1JA Clone C4). Inserts show DAPI staining of cell nuclei. **D:** H1JA and Clone C4 cNES cells could differentiate to TBR1 positive neurons (TUBB3) in ten days. Arrows indicate TBR1 and TUBB3 positive neurons. Scalebar: 20μm **E:** Heatmap of differentially expressed genes among four biological groups of samples. Gene expression levels are presented by Z-score. **F:** MA plot representation of differentially expressed genes between cNES and mhNES clones. Differentially expressed genes are highlighted in light red, selected stem cell (grey), dorsal forebrain (green), ventral forebrain (orange), midbrain (blue) and hindbrain (purple) related genes are named and highlighted with corresponding color. **G:** Schematic representation of main developmental signal transduction pathway components. Coloring indicates Z-score of each gene. Squares indicate average of samples of cNES cell clones (upper), mhNES cell clones (lower).

Next, we analysed the gene expression profile of cNESCs and compared it to that of more posterior brain regions such as midbrain and hindbrain. We found that both bulk cNESC cultures from H1, CA1 hPSCs in 6F (termed H1JA, CA1J) (Figure 2C, S2C) and single cell-derived clones from H1JA that expressed FOXG1, OTX1/2 and PAX6 (Figure 2C, S2C) could differentiate to TBR1 positive neocortical projection neurons (Figure 2D, S2D). In order to determine whether cNESCs have a forebrain-specific gene expression signature, we compared the transcriptome of 6F cNESCs (Group 1; H1JA-derived clones and CA1J and H1AJ-derived bulk cultures); hPSC differentiated to dorsal forebrain neuroepithelium for 8-10d without culture in 6F condition (Group 2); midhindbrain specified NESCs (mhbNESCs) (Group 3; derived in FGF and EGF from hPSCs-AF22, 23 (Falk et al., 2012)- or from human embryos-SAI1,3 (Tailor et al., 2013)-); and hPSCs (Group 4; H1, H7, H9, and CA1), (Figure 2E). 6F cNESCs showed different gene expression signatures from mhbNESCs and hPSCs (Figure 2E) and were more similar to dorsal forebrain specified cultures (Group 2). mhNESCs (Group 3) clustered differently from both hPSCs (Group 4) and all dorsal forebrain specified samples (Group 1) (Figure 2E).

We then analyzed gene clusters that were specifically expressed in cNESCs. Our analysis revealed that 6F cNESCs (Group 1) expressed genes related to neural differentiation, neurogenesis, axon development, cell-cell signalling and cell adhesion. We detected increased expression of genes related to forebrain development, cilium morphogenesis and negative regulation of both neurogenesis and gliogenesis (Cluster E, Figure 2E, S2C). Since we were able to detect the enrichment of forebrain development-specific genes in our transcriptome analysis of 6F cNESC samples, we specifically examined genes related to forebrain and mid-hindbrain specification and development in 6F cNESCs and mhbNESCs (Group 3). Both cNESCs and mhbNESCs had similar expression levels of neural progenitor genes *SOX1, SOX2, NES, NOTCH1, PAX6* (Figure 2F). Strikingly, 6F cNESC clones showed upregulation of forebrain-specific genes, including *OTX2, FOXG1, FEZF2, SP8, EMX2, SIX3* while mhNESCs had showed higher expression of *DPPA4, PAX3, EN2, GBX2, IRX2, HOXB2, GLI2* and *HHIP*, thus demonstrating distinct gene expression patterns that reflect their respective anatomical regions (Supplementary Figure 3A-C).

We further examined the differential expression of genes important for developmental signalling in these two (6F cNESC and mhbNESC) populations (Figure 2G). When we examined TGFβ pathway component genes we found that both *BMPR1B* and *TGFβR1-2-3*, receptors of the TGFβ superfamily were expressed at high levels in mhbNESCs compared to cNESC, while Activin receptor type 1 and 2 (*ACVR1*, *ACVR2*) were expressed at the same level. This matches our observations that cNESCs respond to BMP and ACTIVIN A but not TGFβ1 (Figure 1J). When we examined mediators of the WNT signalling pathway, we found increased expression of a CTNNB1 inhibitor protein gene *ICAT/CTNNBP1* in cNESC cells, which is required for *FOXG1* expression and forebrain specification of the neuroectoderm (Satoh et al., 2004). We detected increased expression of *WNT5a, DKK1* and *DKK2* in mhbNESCs (Figure 2G). However, the increased expression of DKK was not sufficient to reduce CTNNB1 transcriptional activity, as measured by elevated target genes *AXIN2* and *LEF1* levels (Figure 3E). Analysis of FGF and EGF signalling pathway components showed that FGF receptors were expressed at similar levels in both cNESC and mhNESCs, but *FGF2* mRNA levels were increased in cNESCs. In contrast, *EGFR* and *ERBB4* expression was higher in mhbNESCs. When we looked at downstream signalling components we found that the expression of adapter proteins genes (*SHC1, GAB1*), *HRAS, PI3K* genes (*PIK3R1, PIK3R2, PI3KCB*) and *PDK1, AKT1, AKT2* were downregulated in cNESCs and upregulated in mhNESCs. The negative regulator of PI3K (*PTEN*) was higher in cNESCs compared to mhNESCs, while MAPK pathway components (*RAF, MEK, ERK*) were expressed at the same level in both cell types. When we investigated the downstream PI3K and MAPK pathways, we found that cNESCs had reduced expression of several cell cycle regulators like *MYC, ELK1, CDK2* and *CCND2* compared to mhNESCs. We also detected reduced expression of adapter protein 14-3-3 (*YWHAB, YWHAH, YWHAE*) that is important for exporting phosphorylated targets of AKT kinases, such as the Forkhead family proteins from the nucleus. Thus, indicating an overall reduction in the responsiveness to PI3K/AKT activity in 6F cNESCs. Altogether, the differential expression of components of the TGFβ, WNT, MAPK and PI3K pathways in cNESCs compared to mhNESCs, supports our observations that cNESC maintenance preferentially requires regulation of 6F targets than combined activation of FGF and EGF receptors.

**Figure 3.**
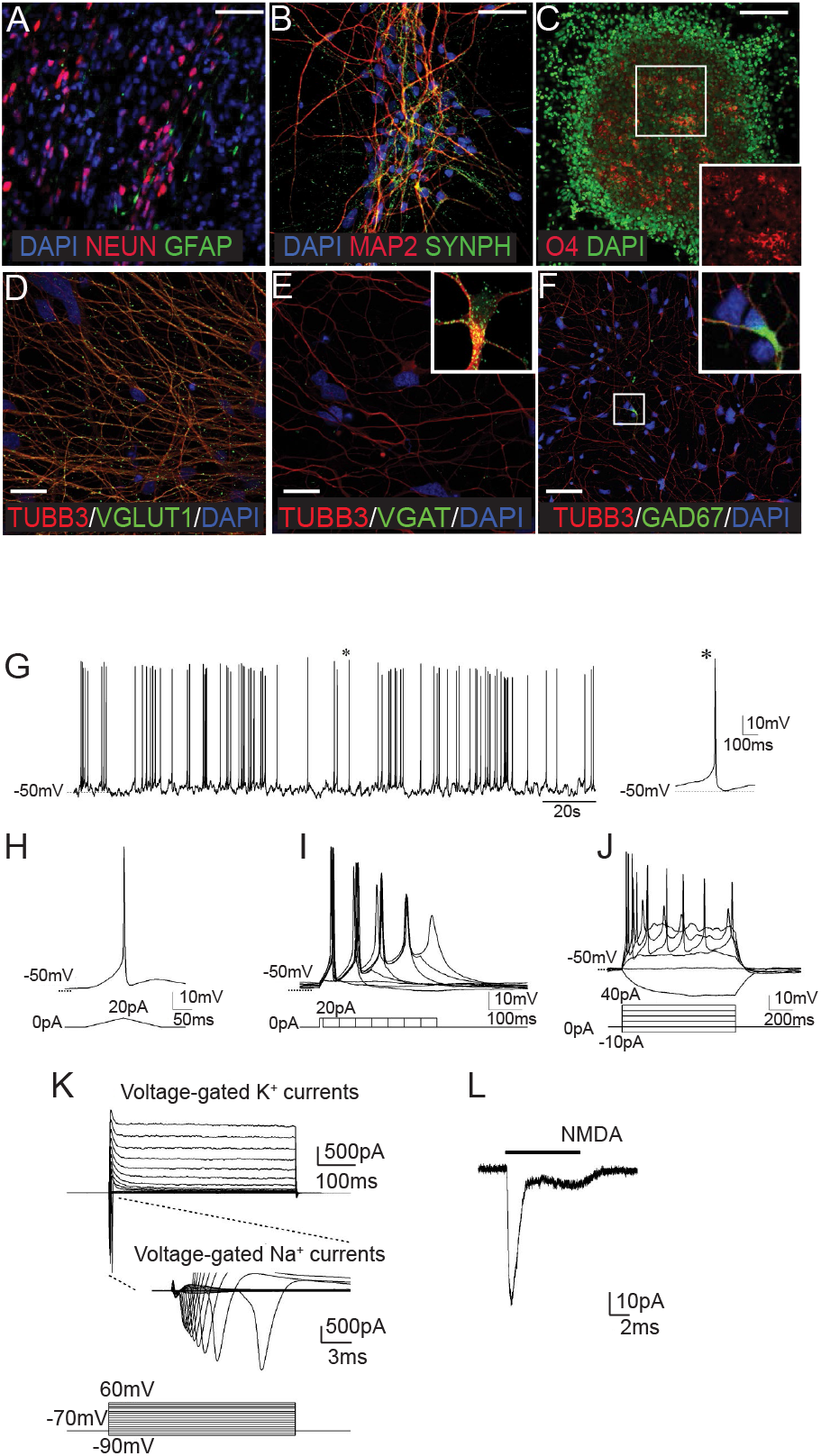
cNES cells can differentiate to mature neurons and glial cells. **A:** cNES (1.53E) cells differentiate to NEUN positive neurons and GFAP positive astrocytes. Scalebar: 20 μm **B:** Mature MAP2 positive neurons express SYNAPTOPHYSIN by day 30. Scalebar: 20μm. **C:** CNES (1.53E) cells differentiate to O4 positive oligodendrocytes, insert shows colony in higher magnification. Scalebar: 50μm. **D:** The majority of cNES (H1A) neurons are glutamatergic projection neurons positive for vesicular glutamate transporter type 1 (VGLUT1). Scalebar: 10 μm **E:** Among the projection neurons (negative for VGAT) a subpopulation of GABAergic interneurons exist, expressing vesicular GABA transporter (VGAT, high magnification from a different area in the insert). Scalebar: 10 μm **F:** GABAergic interneurons also express glutamate amino decarboxylase 67 enzyme (GAD67, the squared area is shown in high magnification in the insert). Scalebar: 20 μm **G:** Representative spontaneous action potentials (APs) in differentiated H1A cNES neurons. A single typical action potential was shown in the right panel. **H:** A single AP was evoked by the ramp current injection (0 to 20 pA). **I:** Trains of APs could be initiated with extend current injection (20 pA, 10 - 360 ms at 50 ms increment). **J:** APs were evoked by step currents injection (−10 to 40 pA). **K:** Voltage-gated K+ and Na+ currents were detected following the depolarizing voltage steps (90 to +60 mV at 10 mV increment). **L:** N-methyl-D-aspartic acid (NMDA)-gated currents could also be evoked in a neuron (hold at - 70mV) by exogenous NMDA (1mM) application.

### cNESC derived cortical neurons are mature and functionally active

Next, we tested whether the differentiation potential of the cNESCs was maintained after prolonged culture. We differentiated cNESCs maintained *in vitro* for 75 days. After 30 days of differentiation, cultures contained β3-tubulin (TUBB3)-positive neurons (approx. 60% of cells) that also expressed MAP2 and NEUN (approx. 40% of cells) (Figure 3A,B), indicating that cNESCs formed mature neurons. Mature MAP2-positive neurons formed presynaptic complexes, as shown by the expression of synaptic protein SYNAPTOPHYSIN (Figure 3B). cNESCs could also differentiate into GFAP positive astrocytes, and O4 positive oligodendrocytes (Figure 3A,C). Thus, confirming that cNESCs maintain multipotency during long-term culture.

To further characterise the types of cortical neurons produced, we analysed their neurotransmitter expression. The majority of cells were glutamatergic projection neurons (vesicular glutamate transporter type 1 (VGLUT1)-positive) (Figure 3D), while GAD67 positive and vGAT positive interneurons comprised around 10% and 2%, respectively, of the total number of neurons (Figure 3E,F).

Next, we determined whether cNESC-derived neurons were mature, electrophysiologically active cells. After 45 days of *in vitro* differentiation, we observed spontaneous action potentials in around 50% of the cells (Figure 3G). These cells could be further evoked to fire action potentials by applying ramp current stimulation (Figure 4H). When constant currents (+20 pA) were held for an extended period (10-360 ms) (Figure 3I) or increasing the current steps from −10 pA to +40 pA up to 1000 ms (Figure 3J), trains of action potentials were seen. With increased depolarizing voltage steps, we were able to detect both fast voltage-gated Na^+^ currents and subsequent voltage-activated K^+^ currents (Figure 3K), confirming the presence of necessary ion channel components for action potentials. Moreover, addition of glutamate receptor agonist *N*-methyl-D-aspartic acid (NMDA, 1 mM) and consequent activation of postsynaptic receptors resulted in the induction of excitatory membrane currents (Figure 3L). Together, these results demonstrate the functional maturation of neurons derived from 6F cNESCs.

**Figure 4.**
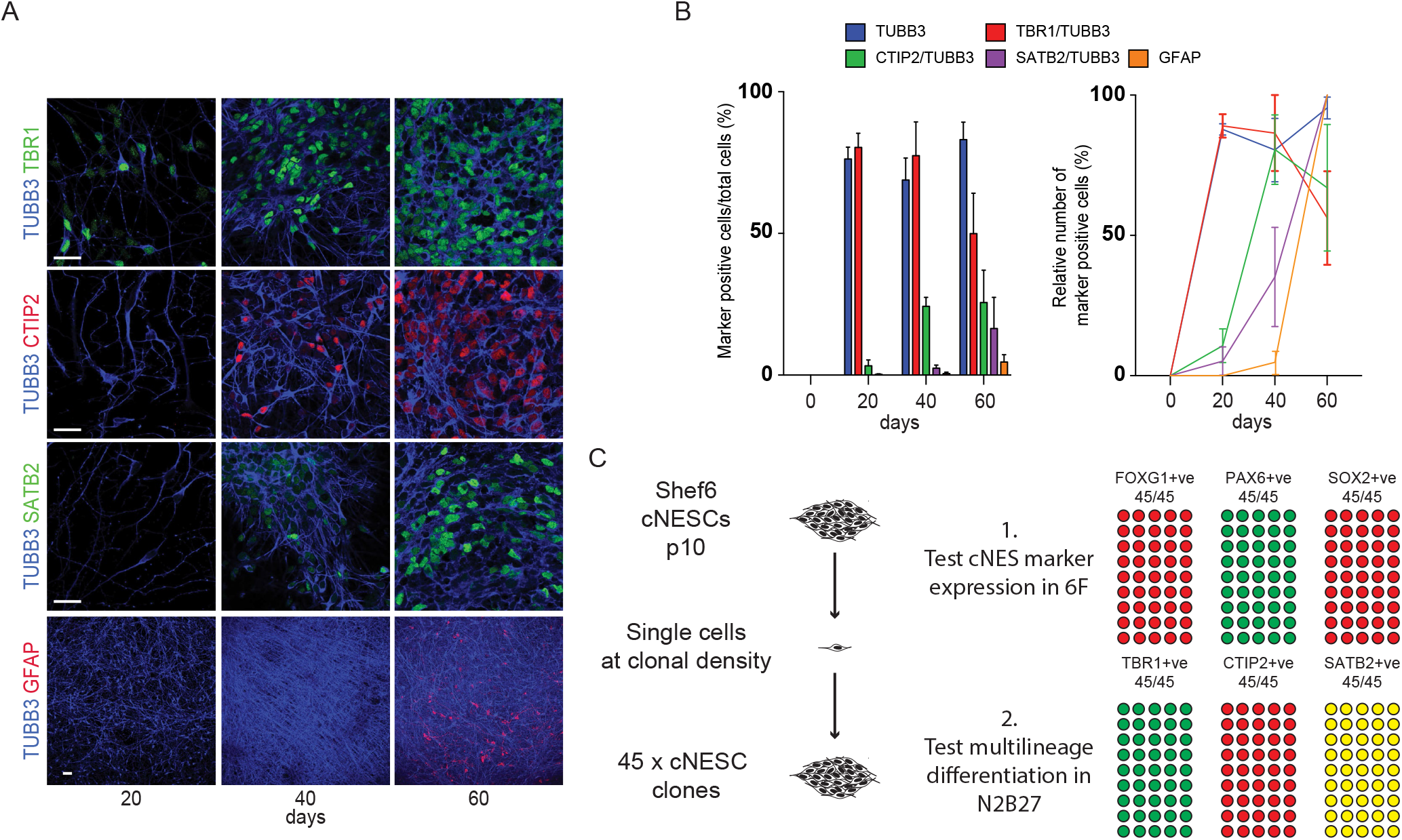
cNES cells in 6F maintain cortical developmental potential. **A:** Cortical layer specific TBR1-CTIP2-SATB2, neuron specific TUBB3 and astrocytic GFAP marker labelling of differentiated SHEF6 derived cNES (passage 12) cells at various time points. Scalebar: 10 μm. **B:** Quantification of cortical layer specific marker positive neurons and astroglial cells in cNESC (passaged for at least 12 times in 6F) differentiating cultures (H1JA, CA1J, S6, H1JA-C4), left panel shows percentage of marker positive cells, right panel shows normalized numbers, 100% is the highest value during the time course (Mean values with SEM, N=4). **C:** Schematics of isolation of clonal cNES cells in 6F condition (left). Results of immunofluorescent labelling of indicated cNES cell markers (top right) and differentiation markers of various cortical layers (bottom right) in 45 clonal cNESC populations. Each circle represents a cNESC clone, filled circles represent marker positive clones. TBR1 and CTIP2 expression was tested at day 30 and SATB2 was tested at day 60 of differentiation.

### 6F recapitulates cortical neuron differentiation timing *in vitro*

During development, the differentiation of glutamatergic projection neurons follows an “inside first, outside last” pattern, which results in the formation of the six layers of the neocortex. To determine whether 6F cNESCs could recapitulate this pattern, we examined the potential and timing of cNESC differentiation to lower and upper layer neurons. Strikingly, we saw the layer 6, TBR1 positive neurons appear first (Figure 4A,B), followed by layer 5 specific CTIP2 positive neurons (Figure 4A,B) and finally the appearance of upper layer specific SATB2 positive neurons (Figure 4A,B). The generation of neurons was followed by the emergence of GFAP positive astrocytes (Figure 4B). This showed that 6F cNESCs preserve the “inside first, outside last” embryonic pattern of differentiation (Figure 4B, right panel).

Next, we tested if individual cNES cells can retain multilayer cortical differentiation potential if maintained in 6F condition. We plated Shef6 (passage 10) cNES cells at clonal density in a 10cm dish in 6F medium and isolated individual colonies (48) after 10 days to subculture them in 96 wells to keep clones for long-term culture (Supplementary Figure 4A). Forty-five out of forty-eight colonies could be plated and expanded in the 96 well plate. Next, we tested 1) if the Shef6 cNESC clones maintained the dorsal forebrain specification in 6F condition by immunofluorescent labelling of FOXG1, PAX6 and SOX2 in the cells and 2) if the clones maintained multilayer specific differentiation potential by differentiating them in N2B27 and labelling deep layer cells with TBR1 and CTIP2 and upper layer cells with SATB2. All cNES clones (45/45) expressed SOX2, FOXG1 and PAX6 markers (Supplementary Figure 4B), demonstrating the maintenance of dorsal forebrain specification in the 6F condition. When we tested if the S6 clones can differentiate into multiple cortical layer specific cells, we found that all clones (45/45) had the potential to differentiate TBR1 positive layer 6 and CTIP2 positive layer 5 cells in 30 days (Supplementary Figure 4B). After sixty days of differentiation we were able to identify SATB2 positive upper layer cells in all differentiating clones (Supplementary Figure 4B) demonstrating the maintenance of multilineage potential of cNES cells in the 6F condition.

### Transplanted cNESCs execute an early developmental cortical programme *in vivo*

We then asked if the early developmental differentiation programme of the cNESCs could be activated *in vivo*. We transplanted undifferentiated, proliferating 6F-cNESCs into the cerebral cortex of 9-week-old juvenile mice (Figure 5A). Seven-weeks post-transplantation, we saw formation of rosette-like polarised structures (indicated by arrowheads) that contained KI67 positive cycling progenitors (Figure 5B) and NESTIN positive cells (Figure 5C). By thirteen weeks cNESCs differentiated into a large number of hNCAM positive neurons (Figure 5D). When we analysed these cells for cortical layer specific markers, we found TBR1 and CTIP2 positive deep layer cells (Figure 5E,F) and CUX1 positive upper layer cells (Figure 5G) in distinct, non-overlapping populations of cells. Both deep and upper layer specific human cells gave rise to layer specific, hNCAM positive neurons (Figure 5H-J). As upper layer neurons in the cortex project to the contralateral hemisphere via the corpus callosum, we looked for the expression of hNCAM in the hemisphere contralateral to the transplant. We noticed the presence of several hNCAM processes and the absence of HUNU positive nuclei in the contralateral corpus callosum, suggesting that cNESC-derived cortical neurons are able to project long distances (Figure 4K). These results show that, not only cNESCs maintain their potential to generate both deep and upper layer neurons *in vitro* but also *in vivo*.

**Figure 5.**
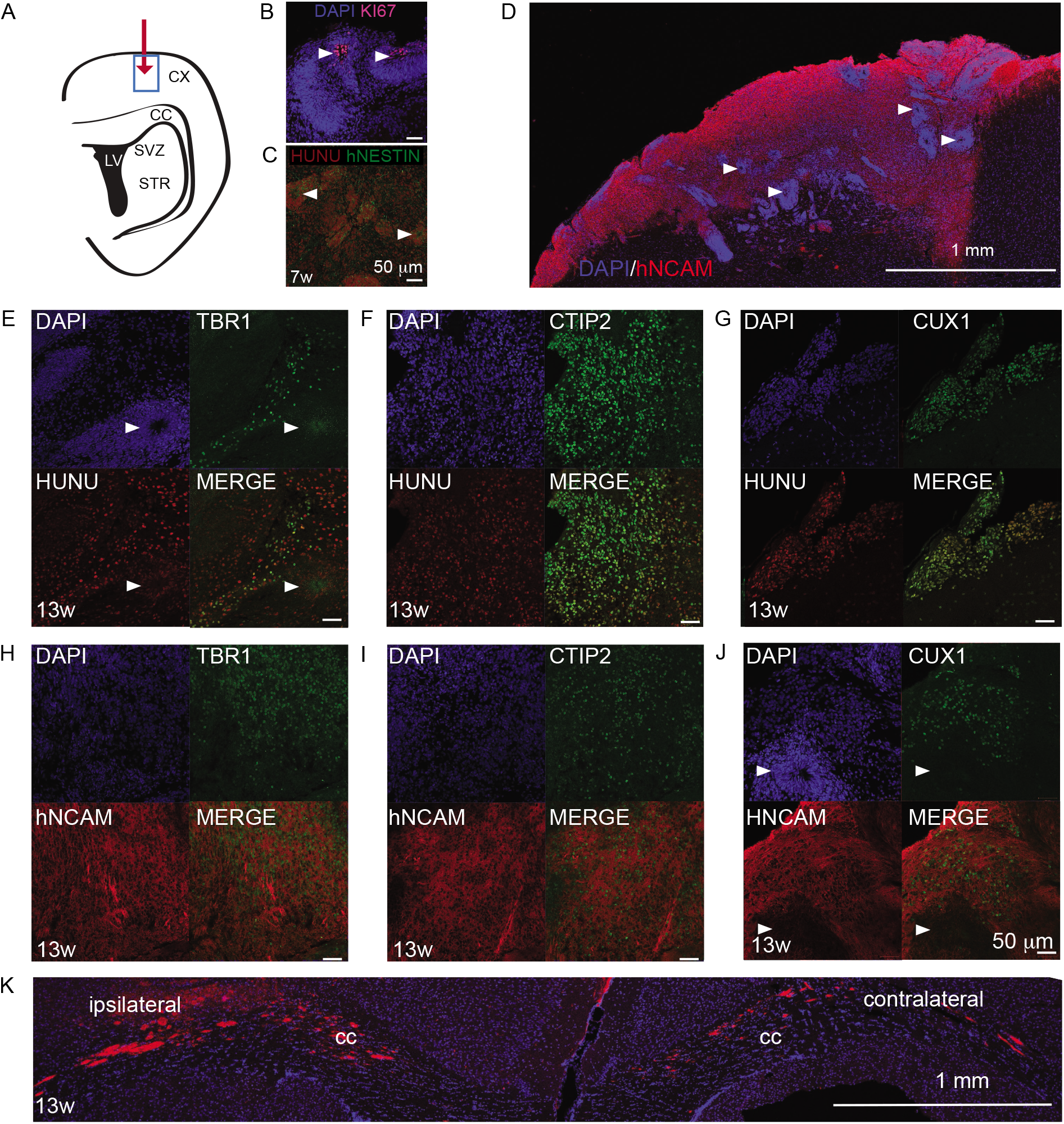
Transplanted cNESCs can complete embryonic cortical differentiation potential in the postnatal brain. **A:** Schematic of 6-week-old mouse coronal brain section indicating the injection site (red arrow) and analysed area (blue rectangles). **B:** After 7 weeks post transplantation polarized rosette structures (arrowheads) of transplanted cNESCs (H1JA, CA1J) contain proliferating KI67 positive human cells. **C:** NESTIN positive human progenitors were found in the grafts at 7 weeks. **D:** Human NCAM expressing neurons are abundant in grafts of CNES cells seven weeks after transplantation. **E-J:** Transplanted human CNES cells differentiated *in situ* to TBR1 (**E, H**), CTIP2 (**F, I**) and CUX1 (**G, J**) positive HuNu and hNCAM expressing cells by 13 weeks after transplantation. **K:** Human NCAM positive neuronal neurites were detected in the contralateral hemisphere of transplanted mice after 13 weeks post transplantation. Arrowheads indicate polarised neural rosettes.

## Discussion

We investigated the role of developmental signalling pathways to determine the factors necessary for maintenance of multipotent cortically-specified neuroepithelial stem cells (cNESCs) *in vitro*. By testing combinations of activators and inhibitors, we found a combination of six regulator factors (activation of FGFR together with inhibition of BMPR, TGFβR, GSK3, AKT and TNKS; termed 6F) that is able to support self-renewal of human cNESCs.

Cortical or dorsal forebrain neuroepithelial cells provide the founder progenitor cells for the development of the cerebral cortex. In the embryo, these cells express markers SOX1, SOX2, FOXG1, OTX2, EMX2, PAX6 and LHX2 and have the potential to generate glutamatergic neurons of the lower (TBR1+, CTIP2+) and upper (BRN2+, CUX1+, SATB2+) layers of the cortex, astrocytes and oligodendrocytes.

Similar to mouse forebrain neuroepithelium (Paek et al., 2009), we found that FGFR activity is needed for human cNES cell proliferation, and this is mediated by both MAPK and PI3K pathway, however AKT inhibition allows colony formation and proliferation. An additional key component of the self-renewal of cNESCs is the regulation of GSK3 and CTNNB1 activity, which is similar to the mouse embryonic forebrain neuroepithelium (Andoniadou et al., 2011; Ferrer-Vaquer et al., 2010; Kim et al., 2009; Masek et al., 2016; Roth et al., 2010; Satoh et al., 2004). Inhibition of GSK3 promotes proliferation and colony formation of human cNESCs but the level of transcriptional CTNNB1 activity regulates the expression of essential dorsal forebrain specification genes FOXG1, EMX2, PAX6, and OTX1/2. Combined inhibition of GSK3 and CTNNB1 transcriptional activity supports cNESC proliferation, colony formation and the maintenance of dorsal forebrain specification and differentiation potential. GSK3 interacts with ERK to downregulate SMAD mediated ACTIVIN A and BMP signalling. Therefore, reduction of GSK3 activity in cNESC requires the inhibition of BMP and ACTIVIN A for the self-renewal of cNESCs. BMP inhibition in cNESCs maintains the expression of SOX1, an early marker of neuroepithelium, while inhibition of ACTIVIN A signalling maintains the proliferation and allows colony-formation of cNESCs.

Our developmental signal screen determined that 6F cNESCs do not require NOTCH activation by gamma secretase-mediated cleavage. NOTCH signalling has been shown to be important in cortical development, but dispensable during the neuroepithelial expansion phase; nonetheless how cNESCs could compensate for a lack of NOTCH activation is not known. Previous studies (Lowell et al., 2006) have indicated that during mouse neural induction *in vitro*, NOTCH signalling is only required for the switch from a neuroepithelial to radial glia-like state. Our findings suggest that differentiation of neural progenitors can be inhibited even if gamma secretase activity mediated NOTCH signalling is reduced.

We also found that 6F cNESCs show a reduced expression of EGFR compared to mhNESCs and do not require EGFR signalling. A study demonstrated that long-term exposure of SHH treated human forebrain neuroepithelial cell to FGF and EGF causes a their transition to radial glia-like cells and differentiation into astrocytes (Edri et al., 2015). EGFR receptor-mediated signalling is known to be important for the gliogenic differentiation of neural progenitors (Aguirre et al., 2010). Moreover, EGF induces elongated bipolar morphology in mouse cortical radial glia cells, allowing the cells to act as scaffold for migratory cells (Gregg and Weiss, 2003). Together, this may explain why previous attempts using a combination of FGF and EGF failed to promote early cNESC maintenance and supported either radial glia-like cells or mhbNESCs. Our gene expression analysis further highlights key differences in the responsiveness of cNESCs to FGF and EGF. EGF receptors and components that regulate the PI3K pathway and HRAS of the MAPK pathway are downregulated while PTEN, the inhibitor of PI3K signalling, is consistently upregulated in cNESCs compared to mhbNES cells. Additionally, we found that cNESCs can be maintained in the absence of SHH signalling and can proliferate and form colonies if SHH, EGF and NOTCH signalling is inhibited.

One of the key properties of self-renewing stem cells is the sustained differentiation potential over rounds of cell divisions. 6F cNESCs retain early cortical potency long-term and preserve the developmental order of corticogenesis. Once the cNESCs are allowed to differentiate by the removal of the 6F neurons of the TBR1 and CTIP2-positive deep layers are produced first, followed by SATB2 and CUX1 positive upper layer neurons and later astrocytes. Although cNESCs express markers of the neuroepithelium (*SOX1, SOX2, HES5* and *PAX6*) in the absence of TNKS and AKT inhibitors (4F), they show reduced expression *FOXG1, OTX2* and *EMX2*. As a result of this altered gene expression network, cNESCs do not differentiate to TBR1 positive layer six neurons but instead only generate CTIP2, BRN2 and SATB2 positive glutamatergic neurons (Table S2).

The 6F condition however supports the efficient formation of single cell derived colonies of SOX2 positive cNES cells that express FOXG1 and PAX6 dorsal forebrain progenitor markers indicating sufficient regulation of key signalling components to inhibit differentiation or regional repatterning. The individual cNES clones could differentiate into deep (TBR1, CTIP2) and upper (SATB2) cortical layer specific cells demonstrating the ability of the early dorsal forebrain neuroepithelial cells to proliferate extensively without changing their differentiation potential.

The lack of regenerative potential in the adult and ageing cerebral cortex is hypothesised to be due to the absence of stem cells with cortical potential and the inability of the tissue to regulate the generation or integration of new projection neurons. The transplantation of undifferentiated cNESCs into the juvenile mouse brain demonstrated that cells with early embryonic developmental potential activate their differentiation programme in a juvenile tissue environment and generate deep and upper layer projection neurons. The *in situ* differentiation of transplanted human cNES cells and the survival of cortical neurons for at least 90 days in the adult brain provides an opportunity to investigate the potential of early developmental stage cells to regenerate or recover functions of the adult cortex and the regulating mechanisms of cell lineage specification.

The use of lineage specified cells from induced pluripotent stem cells carrying genomic variants that predisposes the person to a disease provides the opportunity to correlate medical patient history with histopathology and human disease model system that allows the tracing of changes in cell or tissue physiology both *in vitro* or *in vivo*. However, the refinement of the *in vivo* cellular environment is one of the great obstacles to generate human neurodegenerative and developmental disease models *in vitro* or in xenotransplants. The genetic and epigenetic variability between individuals can introduce experimental roadblocks for the comparison of disease target cells. The establishment of up-scalable tissue specific progenitor cultures provide valuable cell source for the generation of substantial number of comparable functional cells for *in vitro* or transplantation studies.

In summary, we present evidence that the control of six signalling pathway components controls the self-renewal of early cortical neuroepithelial stem cells *in vitro*. In this cell state, single cNESCs can generate populations of identical cells independent of NOTCH, SHH and EGFR signalling. The cNESCs preserve their early developmental potential and neuroepithelial cell state until the 6 signalling components are under control and this allows the transplanted human cNESCs to differentiate *in situ* to projection neurons of the cerebral cortex in the juvenile and adult brain.

## Experimental procedures

### Human pluripotent stem cell culture and neural induction

Human pluripotent stem cells (hPS) were routinely cultured on mitomycin C treated mouse embryonic fibroblast feeder cells in KSR media with 10ng/ml FGF2. KSR media contains 80% DMEM-F12 (GIBCO), 20% Knockout serum replacement (GIBCO), 0.1mM 2-mercaptoethanol, 2mM Glutamax, 0.1mM Non-essential amino acids. Prior to neural differentiation hPSCs were transferred to geltrex (GIBCO) coated surface in the absence of feeders and were cultured in mTESR2 (Stem cell technologies) media. On the day of neural induction, hPSCs were replated on geltrex coated surface in N2B27 media supplemented with 10μM SB431542 and 100nM LDN193189 and 10μM Y27632. For further details please see Supplementary Table 1.

### Neuroepithelial stem cell maintenance and differentiation

Neural cultures were detached from Geltrex first on day 8-10 with Accutase or PBS-EDTA, and replated in neural maintenance media (NES) with 10μM Y27632 at 3×10^5^ cells per cm^2^ on laminin (Sigma) coated surface. NES media contains 50% DMEM-F12, 50% Neurobasal, 0.1mM 2-mercaptoethanol, 2mM Glutamax, 1x N2 supplement, 0.05x B27 minus vitamin A supplement (GIBCO). 4F media is made by supplementing NES media with 10ng/ml FGF2, 3μM CHIR99021, 1μM SB431542 or 50ng/ml FST (Peprotech), 100nM LDN193189 or 50 ng/ml NOGGIN. 6F media contains components of the 4F media with 100nM K02288 (Selleckchem), 100nM AKTiVIII and 75nM MK2206 (Selleckchem), and 1-2 μM XAV939 (Selleckchem). XAV939 concentrations need to be adjusted to each cell line, 1μM for H1, CA1, H9, CTRL hPSCd-erived cNESCs, 2 μM for Shef6 derived cNESCs. For oligodendrocyte differentiation cNESCs were cultured in 4F and cells were collected in clumps using EDTA-PBS. Floating cell clumps were cultured in 4F media for 2-3 days and the aggregates were transferred to Neural Differentiation I Media. Aggregates were cultured for 30 days in Neural Differentiation I Media with media changes every 4 days. Floating aggregates were plated onto Poly-D-lysin and laminin coated glass coverslips and cultured for 14 more days in Neural Differentiation I Media. Cultures were fixed with 4% Paraformaldehyde and stained with anti-O4 mouse IgM antibody. For further details please see Supplementary Table 1.

### Proliferation assay

Proliferation of hNESCs cells was assayed by plating 1×10^4^ cells per each laminin coated well of a 96 well plate in NES media. Cells were allowed to attach and spread for 2 hours and one volume of seeding media was complemented with equal volume of test media with 2x concentration test molecules. Tested molecules were used in dilution series and had the following highest final concentrations: ACTIVIN A (100ng/ml, Peprotech), BMP4 (20ng/ml, Peprotech) FST (50ng/ml, Peprotech), NOGGIN (50 ng/ml, Peprotech), CHIR99021 (3μM, Selleckchem), Cyclopamine-KAAD (200nM, Millipore), DAPT (5μM), EGF (100 ng/ml, Peprotech), FGF2 (10 ng/ml, Peprotech), LDN193189 (100nM, Selleckchem), PD0325901 (1μM, Selleckchem), PD153035 (2.5μM), PD173074 (0.3μM), PI-103 (0.5μM), Wortmannin (0.1μM), Purmorphamine (1μM), SAG (100nM), SB202190 (20μM), SB203580 (20μM), SB505124 (1μM), SB431542 (10μM), LY364947 (5μM), TGFβ1 (100ng/ml), Y27632 (10μM), JNK-IN-8 (3μM), SP600125 (1μM). Cells were incubated with the indicated factor for 96 hours. Total cell numbers were determined by Alamar blue cell viability assay (Life Sciences) based on cell number titration. Each value is the average of 6 technical replicates, average values were normalised to control condition (4F) and represented as relative values. All experiments were repeated using three cell lines (H1A, H9A, H7E).

### Clonogenicity assay and single cell clone isolation

H1 (passage 12) and S6 (passage 9) cNES cells were cultured in 4F or 6F medium as described in section ‘Neuroepithelial stem cell maintenance and differentiation’. Three to four days after plating the cells were rinsed with PBS once and detached with 5-minute Accutase digestion at 37°C. The cells were collected with 10fold volume DMEM-F12 with 0.1% BSA and centrifuged at 300xg for 3.5 minutes. The pelleted cells were resuspended in 1 ml fresh 6F media and triturated by pipetting with 1000μl tip for 20 times with moderate force without generating bubbles. Cells were centrifuged with additional 5fold volume DMEM-F12 with 0.1% BSA at 300xg for 3.5 minutes. For clonogenicity assay the pelleted cells were resuspended in 1 ml fresh NES media with FGF2 at 20ng/ml concentration and counted in a haemocytometer. Two hundred cells per cm^2^ culture surface area were plated in poly-D-lysine and laminin coated plastic culture dishes. Two hours after the plating cells attached and spreaded well, and media was changed to test conditions. Fresh FGF was added daily at 20ng/ml concentration without media change for the first 4 days until clones reached a colony size of 4 or more cells. Media was completely changed after 6 days.

Cultures were fixed at indicated timepoints with 4% PFA solution for 10 minutes at room temperature. After rinsing with PBS the cell colonies were labelled with Cresyl violet solution for 5 minutes at room temperature and rinsed with water. The plates were scanned and colonies were counted with Volocity software. Occasionally colonies split to form 2-3 small colonies close to each other and separate from neighbouring colonies. The split colony clusters were counted as one. For single cell cloning S6 cNES cells were plated in 6F media at 100 cells per cm2 culture surface area in a 10 cm dish coated with poly-D-lysine and laminin. FGF was added daily at 20ng/ml concentration without media change for the first 4 days. Media was completely changed after 6 days. Colonies were manually picked at day 10 to 96 wells. Clonal S6 cNES cells were maintained in 96 wells and passaged once 1:4 ratio for immunofluorescent labelling of cNES cells or *in vitro* differentiation assay. Cultures in 6F were fixed with 4% PFA solution for 10 minutes at room temperature.

During the *in vitro* differentiation assay at day 6 the wells were treated with accutase for 2 minutes, accutase was removed and 3 minutes later the cells were gently triturated with N2B27 to generate small clumps of cells. Cells were replated at 1:4 ratio in fresh 96 wells coated with poly-D-lysine and laminin. Cultures were fed every 4 days with 50% of the media volume replaced with fresh N2B27 media that contained 10μM Forskolin and 20ng/ml BDNF. Cultures were fixed at day 30 and 60 with 4% PFA solution for 10 minutes at room temperature.

### Immunocytochemistry

Cultured cells were rinsed once with phosphate buffered saline and fixed with 4% PFA solution at room temperature for 15 minutes. For cell surface epitopes, the cells were stained without Triton X100 in PBS with 5 % fetal bovine serum. Intracellular epitopes were stained after 30 minutes permeabilisation with 0.3% Triton X100 in PBS with 5% fetal bovine serum and 1% BSA. Volocity demo v6.1.1 (Perkin Elmer) was used to quantify nuclear and cytoplasmic immunofluorescent stainings. Staining threshold was calculated from immunofluroescent staining of “epitope negative” cells or secondary antibody controls. Pictures were taken from minimum 6 randomly picked areas.

### Western blot

Cells for western blot experiments were replated the night before collection at sub confluent density, starved for 2 hours for investigated cytokine, rinsed once with DMEM-F12 before treatment. Cells were lysed on ice with RIPA buffer containing protease and phosphatase inhibitors (Complete and PhosSTOP, Roche). Samples were centrifuged for 3 minutes at 15000xg to remove cell debris and treated with DNAse to degrade genomic DNA. Protein concentration was determined by BCA reaction (Thermo) according to the manufacturer’s instructions. Equal amounts of protein (15-50μg) were loaded in equal volumes to SDS-Polyacrylamide gradient gels. Proteins were transferred to nitrocellulose membranes and probed with indicated antibodies. The same membrane was stripped and probed with various antibodies unless otherwise stated.

### Cell transplantation and immunohistochemistry

*In vivo* differentiation of undifferentiated hNESCs cells was tested in NOD-SCID-Gamma adult mice (minimum 6 weeks old). Experiments were carried out with the approval of Mount Sinai Hospital’s Research Ethical Board (REB Approval #12-0106-E) and the Stem Cell Oversight Committee of CIHR. 10^5^ hNESCs cells (H1JA or CA1J) maintained in 6F media was collected with Accutase, centrifuged and resuspended in 6F media. After cell counting, cells were centrifuged and resuspended in DMEM-F12 with 10uM Y27632 at 10^5^ cells per μl and kept on ice for maximum 2 hours before injected into the wall of the lateral ventricle. A 26 gauge Hamilton syringe with a 45 degree bevelled tip was used to stereotaxically inject 1 μl of cell suspension at a rate of 0.1μl/min at the coordinates: 0 AP, 1.4 ML, −1.7 DV, relative to bregma. The needle was left in place for 10 minutes after cell injection to prevent backflow and then slowly withdrawn. Mice were kept individually after surgery and monitored daily until sacrificed for tissue processing. Mice were anesthetized with avertin, perfused transcardially and post-fixed for 2 hours with 4% paraformaldehyde. Brains were cryopreserved in 30% sucrose, frozen and 18 μm horizontal cryostat sections were cut. Cryosections were blocked with 10% normal goat serum (NGS), 0.5% BSA and 0.3% triton and transplanted human cells were labelled with a primary antibody in PBS overnight at 4C, followed by an incubation with a secondary antibody and DAPI (5μg/mL) in PBS for 1h at RT. For double immunohistochemistry, sections were reblocked in 10% NGS, 0.5% BSA and 0.3% triton, incubated with a primary antibody 4°C O/N and then with a secondary antibody at RT for 1h.

### Electrophysiology

Recording of action potentials and ionic currents of the iPS-derived neurons were carried out using standard whole-cell current- and voltage-clamp techniques (Richmond and Jorgensen, 1999). Cells were patched using fire-polished 3–7 MΩ resistant borosilicate pipettes (World Precision Instruments, USA) and membrane currents and voltages were recorded in the whole-cell conFIGuration by a Digidata 1440A and a MultiClamp 700A amplifier, using the Clampex 10 software and processed with Clampfit 10 (Axon Instruments, Molecular Devices, USA). Data were digitized at 10-20 kHz and filtered at 2.6 kHz. To record spontaneous action potentials, the input current was held at 0 pA, and hyperpolarizing and depolarizing step or ramp currents were injected to elicit action potentials. To record voltage-gated K^+^ and Na^+^ currents, cells were held at −70mV, and voltage steps from −90 mV to +60 mV were delivered at 10-mV increments. The pipette solution contains (in mM): K-gluconate 115; KCl 25; CaCl_2_ 0.1; MgCl_2_ 5; BAPTA 1; HEPES 10; Na_2_ATP 5; Na_2_GTP 0.5; cAMP 0.5; cGMP 0.5, pH7.2 with KOH, ~320 mOsm. The external solution consists of (in mM): NaCl 150; KCl 5; CaCl_2_ 2; MgCl_2_ 1; glucose 10; HEPES 10, pH7.2-7.3 with NaOH, ~320-325 mOsm. Leak currents were not subtracted. All chemicals were from Sigma. Experiments were performed at room temperatures (20–22°C).

### Q-PCR

Total RNA was isolated from cultured cells with Qiagen RNeasy kit according to manufacturer’s instructions. One ug of total RNA was reverse transcribed to cDNA with Qiagen reverse transcription kit. Quantitative RT-PCR detection of specific cDNA sequences was done with SYBR Green mix on a BIORAD thermocycler. Primer pairs for each cDNA target were tested for linear amplification and single product melting peak. Ct values are the average of 3 technical replicates, cDNA abundance is normalised to *GAPDH* housekeeping gene reads.

### Microarray analysis

Total RNA was assessed for quality and quantity on a Bio-analyzer and global gene expression profiling performed with the Affymetrix microarray. Purified and labelled RNA was analyzed on Affymetrix Human Transcriptome Array 2.0 (Affymetrix) according to the manufacturer’s instructions, at Microarray Facility of The Centre for Applied Genomics at Sickkids, Toronto, Ontario.

RMA method normalization, background substraction, and summarization was performed using Oligo (v1.34.2) (Irizarry et al., 2003) R package from bioconductor. Normalized gene expression intensities were log2-scaled for all subsequent analyses. Unless otherwise stated, all data presented are representative of at least two independent experiments. A linear model approach and the empirical Bayes statistics were implemented for differential gene expression analysis, as described in limma (v3.26.9) (Phipson et al., 2016) R package user guide. P-values were adjusted using the Benjamini-Hochberg method and significance cut-off was set at 0.01. Euclidean distance matrix and complete linkage were used for hierarchical clustering of the samples using log2 transformed gene expression values. Euclidean distance matrix and Ward’s minimum variance method of clustering were used to obtain the gene clusters using Z-score normalized gene expression values. All statistical analyses, Gene Ontology term and KEGG pathway analyses, and data visualization were done in R using R basic functions and the following packages: clusterProfiler (v2.4.3) (Yu et al., 2012), gplots (v3.0.1, https://CRAN.R-project.org/package=gplots), pathview (v1.10.1) (Luo and Brouwer, 2013) and stats (v3.2.2, http://www.rdocumentation.org/badges/version/stats).

## Authors’ contribution

BVV designed and carried out the experiments, analysed and interpreted the data and wrote the manuscript, MF carried out the transplantation studies and analysed the data, HY and EL carried-out experiments, SG did the electrophysiological recordings and interpreted the data, MZ interpreted the data, SH analysed the microarray data, AN interpreted the data and wrote the manuscript.

## Acknowledgement

The authors would like to thank Ragnhildur Thora Karadottir for help with electrophysiological patch-clamp recordings, Robert Hevner for antibodies, Maria Mileikovskaia for assistance with CA1 hESCs, Gordon Keller for H1, H7, H9 hESCs, Austin Smith for CB660 cell line, Ludovic Vallier for CTRL hiPSCs, Peter W Andrews for SHEF6 hESCs, Chi-chung Hui for Q-PCR primers, Faustine Massin for DNA cloning, Chen He and Puzheng Zhang for technical assistance, Carla Mulas, Masaki Kinoshita, Ian Rogers, Natalie Payne and Kathryn Davidson for critical reading of the manuscript.

---

**Supplementary Figure 1.**
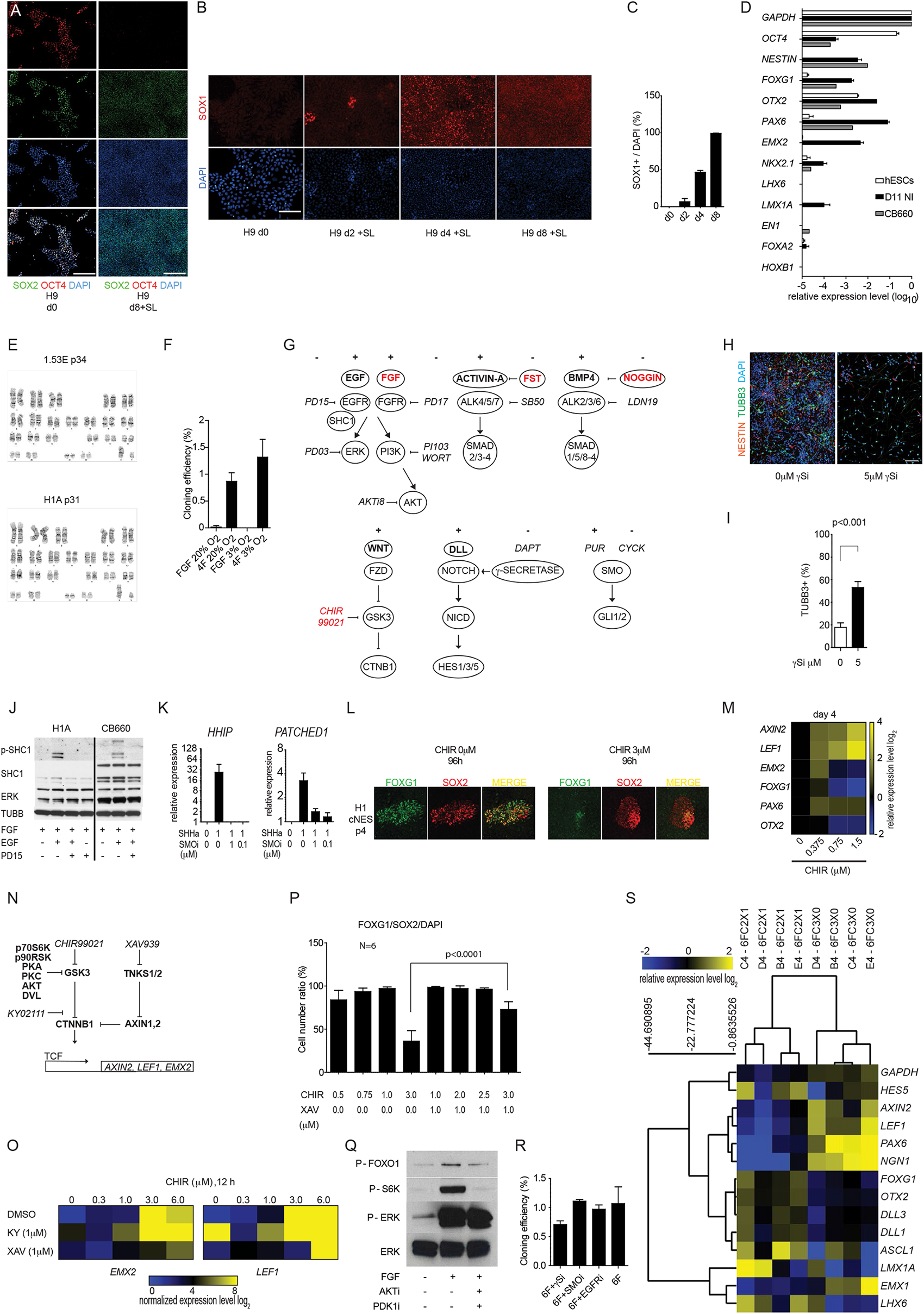
**A:** Monolayer neural differentiation of H9 hES cells in the presence of SB431542 and LDN193189 (SL) reduced the expression of OCT4 and maintained the expression of SOX2 in day 8 cultures. Scalebar:100 μm. **B:** The number of neuroepithelium specific SOX1 positive cells increases over 8 days of neural differentiation of H9 hES cells. Scalebar: 50 μm. **C:** Quantification of SOX1 positive cells during H9 hES cell neural induction, minimum 640 cell nuclei were analysed in 3 technical replicates per time point. **D:** Q-RT-PCR analysis of hPS cells (H1, H9, CA1), corresponding day11 neural cultures and CB660 NS cells. Mean values are relative to GAPDH. **E:** Karyotyping of 1.53E cNES and H1A cells maintained in 4F media. **F:** Colony numbers were counted 11 days after plating. (N=3, mean±SD) **G:** Schematic presentation of developmental signaling pathway components targeted in our assay, protein ligands are in bold and chemical inhibitors are in italic, 4F components are in red. **H:** Inhibition of NOTCH signalling with DAPT induced rapid cell cycle exit and neuronal differentiation of cNES cells (H1A) in 11 days in the absence of 4 factors. Immunofluorescent staining of TUBB3 (green) positive post mitotic neurons and NESTIN (red) positive neural progenitors in day 10 cultures, scalebar 50μm. **I:** Quantification of TUBB3 and DAPI positive cell population. (N=6, Student’s T-test, p<0.001). **J:** Western blot detection of EGFR activity by SHC1 adaptor protein phosphorylation in control (no EGF), EGFR inhibitor PD15 (PD153035, 2.5μM) treated, EGF treated and EGF plus EGFR inhibitor PD15 (PD153035, 2.5μM) treated H1A cNES cells and CB660 NS cells. **K:** Hedgehog signaling can be activated in cNES cells (H1A) by SMOOTHENED agonist Purmorphamine (PUR, 1 μM) and inhibited by the antagonist Cyclopamine-KAAD (CycK). Activation of SMOOTHENED by PUR upregulates mRNA levels of downstream target genes HHIP and PATCHED1 (N=3). **L:** Immunofluorescent staining of FOXG1 and SOX2 proteins in passage four cNESCs (H1 PSC derived) after 96 hour treatment with 3μM GSK3 inhibitor (CHIR) or DMSO. **M:** Q-RT-PCR comparison of mRNA levels of selected dorsal forebrain specific genes and direct CTNNB1 target genes *AXIN2* and *LEF1* after 96hours treatment of H1 cNES cells with various concentrations of GSK3 inhibitor (CHIR) in 4F media, normalised to 0μM treatment. (Values are means of 3 biological replicates). **N:** Schematic of protein and chemical regulators of GSK3 and CTNNB1. Proteins are in bold, chemicals are in italic. **O:** Q-RT-PCR comparison of mRNA levels of EMX2 and LEF1 after 12-hour treatment of p0 H1 cNES cells with increasing concentration of GSK3 inhibitor (CHIR) combined with either XAV939 (XAV) or KY02111 (KY). (N=3) **P:** CNES cells (H1JA) were treated with increasing concentrations of GSK3 inhibitor (CHIR) and XAV939 (XAV) for 96 hours. FOXG1 and SOX2 double positive cell nuclei were counted (N=6, One way ANOVA, Tukey post test). **Q:** FGF activates both MAPK and PI3K/AKT signaling downstream of FGFR. FGF was withdrawn from CNES cells (H1JA) for 2 hours, cultures were incubated for 10 minutes with inhibitors of AKT activity (AKTi, PDKi) before administration of FGF. Reduction of both phosphorylated FOXO1 and S6K levels after FGF treatment but not phosphorylated ERK levels indicates inhibitor treatment effectively reduced AKT activity in cNES cells. **R:** CNES (H1JA) cells were cultured in 6F media and plated at clonal density (200cells/cm2). Addition of NOTCH inhibitor DAPT, SMO inhibitor CYCK or EGFR inhibitor PD15 did not inhibit colony formation of cNES cells (N=3). **S:** Normalized mRNA levels of genes after ranking by significance of changing after withdrawal of TANKYRASE inhibitor XAV939 from 6F media in single cell clones of H1JA cNES cells. Colour bar shows log_2_ level of change from mean value.

**Supplementary Figure 2.**
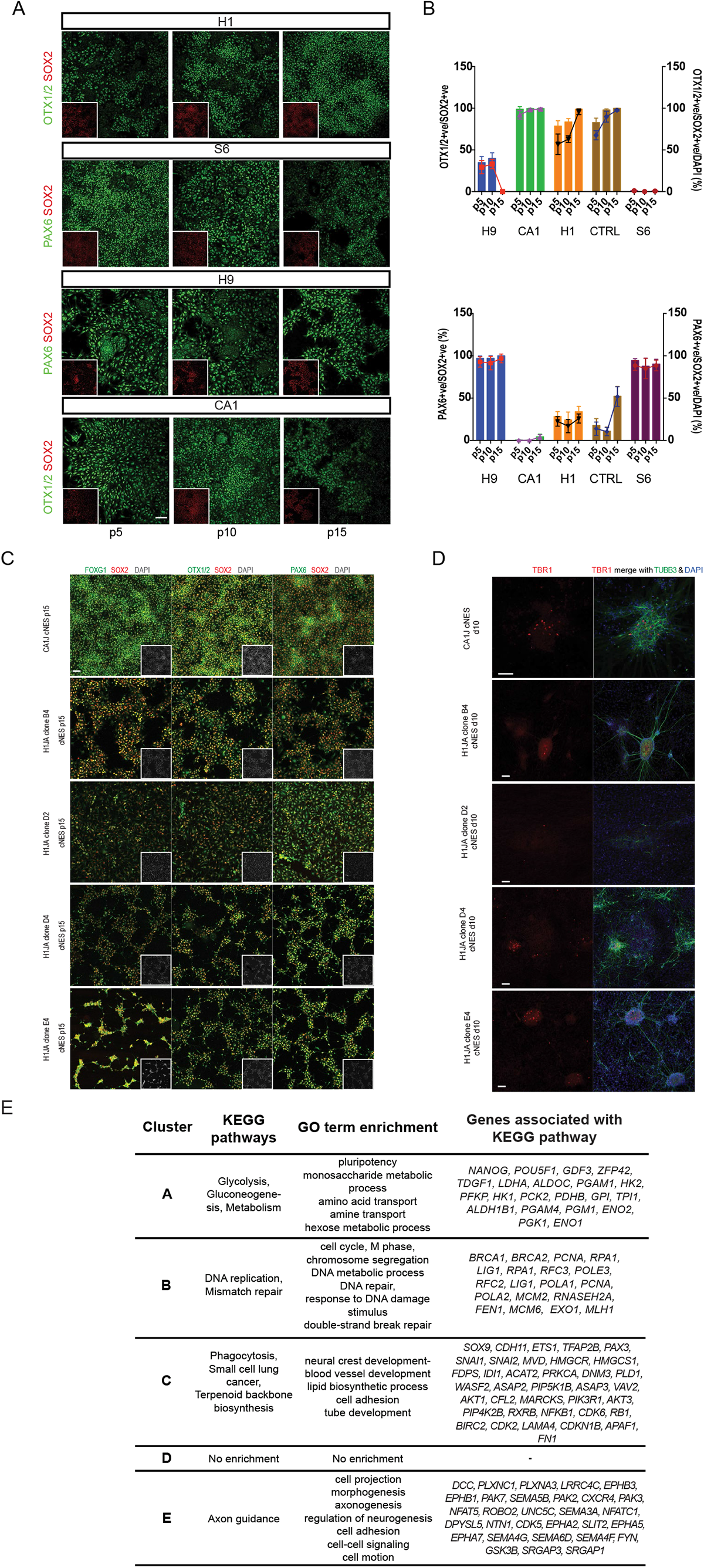
**A:** CNES cells from 4 independent human PSC line (CA1, H1, S6, H9) were cultured in 6F for 15 passages. Immunofluorescent detection of PAX6 or OTX1/2 in SOX2 positive CNES cells at indicated passages. Scalebar: 20μm. **B:** Quantification of OTX1/2, PAX6 and SOX2 positive cNES cells in five independent hPSC derived cNES cell lines from multiple passages. Bars show percentage of OTX2 or PAX6 positive NES cells, lines show percentage of OTX2 or PAX6 and SOX2 double positive cells of all cells. (Mean±SD) **C:** Immunofluorescent detection of cNES markers FOXG1, OTX1/2, PAX6 and SOX2. Inserts show cell nuclei stained with DAPI. Scalebar: 20μm. **D:** Differentiation of cNES cells for 10 days with 5μM DAPT. Immunofluorescent detection of TBR1 and TUBB3 positive neurons in cNESCs. Scalebar: 50μm. **E:** KEGG pathway and GO-term enrichment of differentially expressed genes in each cluster of genes from Figure 2G.

**Supplementary Figure 3.**
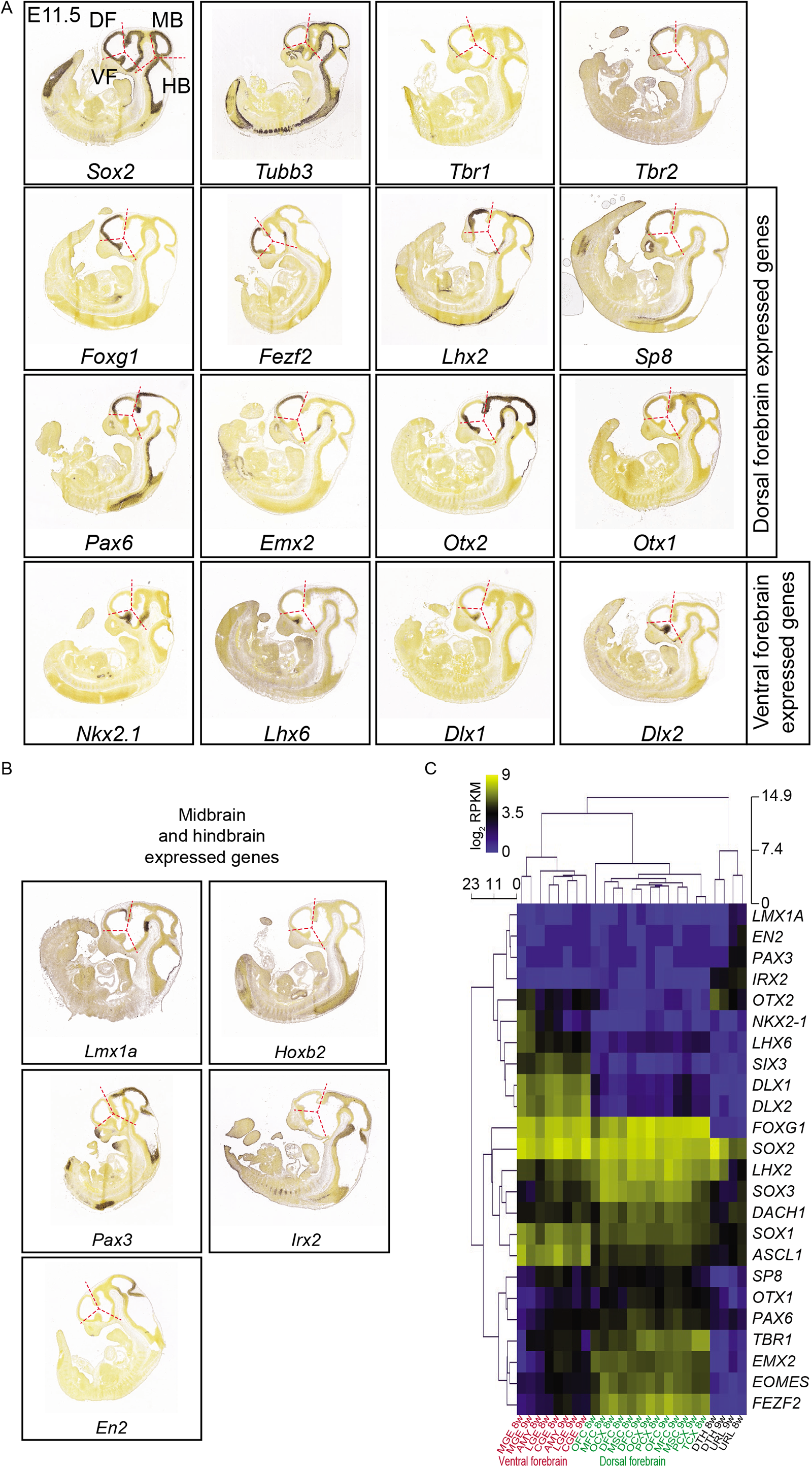
**A:** Expression pattern of dorsal and ventral forebrain marker genes from Figure 2F in the E11.5 mouse embryo sagittal sections. Images are from the Allen Developing Mouse Brain Atlas http://developingmouse.brain-map.org/. The developmental age correlates with the differentiation of TBR1 positive deep layer neurons. **B:** Expression pattern of midbrain and hindbrain marker genes from Figure 2F in the E11.5 mouse embryo sagittal sections. **C:** Expression pattern of cortical layer specific markers from Figure 2C, 4A-C in E18.5 mouse embryo sagittal sections. **D:** RNA-seq expression values of region-specific marker genes in the developing 8-9 week old human brain. Normalised RPKM values are from the BrainSpan Atlas of the Developing Human Brain (http://www.brainspan.org/). MGE-medial ganglionic eminence, LGE-lateral ganglionic eminence, CGE-caudal ganglionic eminence, AMY-amygdaloid complex, OFC-orbital frontal cortex, MFC-mediofrontal cortex, OCX-occipital cortex, DFC-dorsolateral prefrontal cortex, MSC-primary motor and sensory cortex, PCX-parietal neocortex, TCX-temporal neocortex, DTH-dorsal thalamus, URL-upper rhombic lip.

**Supplementary Figure 4.**
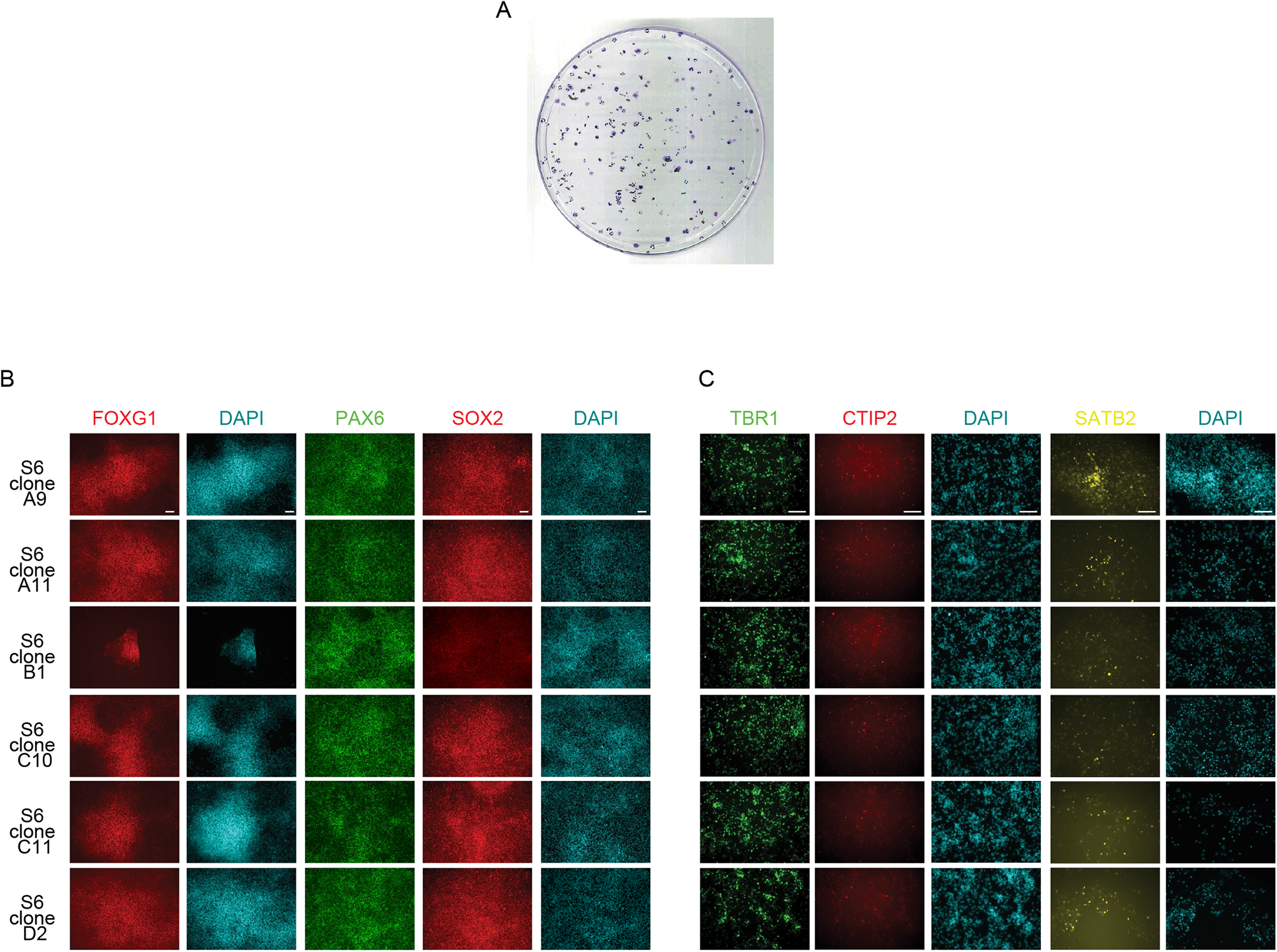
**A:** Cresyl violet staining of day 12 colonies from S6 passage 9 cNES cells maintained in 6F condition. **B:** Examples of immunofluorescent labelling of single cell derived S6 cNES clones. CNES cells were maintained in 6F condition in 96-wells and labelled for FOXG1, PAX6 and SOX2 dorsal forebrain NES cell markers. Cell nuclei were labelled with DAPI. Scalebar: 20μm. **C:** Examples of immunofluorescent labelling of deep layer (TBR1, CTIP2) at day 30 and upper layer (SATB2) at day 60 differentiated cultures of S6 cNES clones in panel B. Cell nuclei were labelled with DAPI. Scalebar: 20μm.

